# Crowdsourcing Image Analysis for Plant Phenomics to Generate Ground Truth Data for Machine Learning

**DOI:** 10.1101/265918

**Authors:** Naihui Zhou, Zachary D Siegel, Scott Zarecor, Nigel Lee, Darwin A Campbell, Carson M Andorf, Dan Nettleton, Carolyn J Lawrence-Dill, Baskar Ganapathysubramanian, Jonathan W Kelly, Iddo Friedberg

## Abstract

The accuracy of machine learning tasks critically depends on high quality ground truth data. Therefore, in many cases, producing good ground truth data typically involves trained professionals; however, this can be costly in time, effort, and money. Here we explore the use of crowdsourcing to generate a large number of training data of good quality. We explore an image analysis task involving the segmentation of corn tassels from images taken in a field setting. We investigate the accuracy, speed and other quality metrics when this task is performed by students for academic credit, Amazon MTurk workers, and Master Amazon MTurk workers. We conclude that the Amazon MTurk and Master Mturk workers perform significantly better than the for-credit students, but with no significant difference between the two MTurk worker types. Furthermore, the quality of the segmentation produced by Amazon MTurk workers rivals that of an expert worker. We provide best practices to assess the quality of ground truth data, and to compare data quality produced by different sources. We conclude that properly managed crowdsourcing can be used to establish large volumes of viable ground truth data at a low cost and high quality, especially in the context of high throughput plant phenotyping. We also provide several metrics for assessing the quality of the generated datasets.

**Author Summary:** Food security is a growing global concern. Farmers, plant breeders, and geneticists are hastening to address the challenges presented to agriculture by climate change, dwindling arable land, and population growth. Scientists in the field of plant phenomics are using satellite and drone images to understand how crops respond to a changing environment and to combine genetics and environmental measures to maximize crop growth efficiency. However, the terabytes of image data require new computational methods to extract useful information. Machine learning algorithms are effective in recognizing select parts of images, butthey require high quality data curated by people to train them, a process that can be laborious and costly. We examined how well crowdsourcing works in providing training data for plant phenomics, specifically, segmenting a corn tassel – the male flower of the corn plant – from the often-cluttered images of a cornfield. We provided images to students, and to Amazon MTurkers, the latter being an on-demand workforce brokered by Amazon.com and paid on a task-by-task basis. We report on best practices in crowdsourcing image labeling for phenomics, and compare the different groups on measures such as fatigue and accuracy over time. We find that crowdsourcing is a good way of generating quality labeled data, rivaling that of experts.

## Introduction

Crop genetics include basic research (what does this gene do?) and efforts to affect agricultural improvement (can I improve this trait?). Geneticists are primarily concerned with the former and plant breeders are concerned with the latter. A major difference in the perspectives between these groups is their interest in learning which genes underlie a trait of interest: whereas geneticists are generally interested in what genes do, breeders can treat the underlying genetics as opaque, selecting for useful traits by tracking molecular markers, or directly, via phenotypic selection (1).

Historically, the connections between plant genotype and phenotype were investigated through forward genetics approaches, which involve identifying a trait of interest, then carrying out experiments to identify which gene is responsible for that trait. With the advent of convenient mutagens, molecular genetics, bioinformatics, and high-performance computing, researchers were able to associate genotypes with phenotypes more easily via a reverse genetics approach: mutate genes, sequence them, then look for an associated phenotype.

However, the pursuit of forward genetics approaches is back on the table, given the even more recent availability of inexpensive image data collection and storage coupled with computational image processing and analysis. In addition, the potential for breeders to computationally analyze phenotypes is enabled, thus allowing for the scope and scale of breeding gains to be driven by computational power. While high-throughput collection of forward genetic data is now feasible, we must now enable the *analysis* of phenotypic data in a high-throughput way. The first step in such analysis is to identify regions of interest as well as quantitative phenotypic traits from the images collected. Tang *et al.* (25) described a model to extract tassel out of one single corn plant photo through color segmentation. However, when images are taken under field conditions, classifying images using the same processing algorithm can yield sub-optimal results. Changes in illumination, perspective, or shading, as well as occlusion, debris, precipitation, and vibration of the imaging equipment can all result in large fluctuations in image quality and information content.Machine learning (ML) methods have shown exceptional promise in extracting information from such noisy and unstructured image data. Kurtulmus and Kavdir (13) adopted a machine learning classifier, support vector machine (SVM), to identify tassel regions based on the binarization of color images. An increasing number of methods from the field of computer vision are recruited to extract phenotypic traits from field data (22, 29). For example, fine-grained algorithms have been developed to not only identify tassel regions, but also identify tassel traits such as total tassel number, tassel length, width, etc. (16, 27)

A necessary requirement for training ML models is the availability of labeled data. Labeled data consist of a large set of representative images with the desired features labeled or highlighted. A large and accurate labeled data set, the *ground truth,* is required for training the algorithm. The focus of this project is the identification of corn tassels, in images acquired in the field. For this task, the labeling process includes defining a minimum rectangular bounding box around the tassel. While seemingly simple, drawing a bound-ing box does requires effort to ensure accuracy (24), and a good deal of time to generate a sufficiently large training set. Preparing such a dataset by a single user can be laborious and time consuming. To ensure accuracy, such a generated set should ideally be proofed by several people, adding more time, labor, and expense to the task.

One solution to the problem is to take a large cohort of untrained individuals to perform the task, and to compile and extract some plurality or majority of their answers as a training set. This approach, also known as crowdsourcing, has been used successfully many times to provide image-based information in diverse fields including astronomy, zoology, computational chemistry, and biomedicine, among others (2– 5, 7, 12, 17, 21, 26).

Crop genetics research has a long history of “low-tech” crowdsourcing. Groups of student workers are sent into fields to identify phenotypes of interest, with the rates of success often a single instance among thousands of plants. Students also regularly participate in experiments to learn about the research process and gain first-hand experience acting as participants. To manage these large university participant pools, cloud based software, such as the Sona system^1^, are routinely used to schedule experiment appointments and to link to web-based research materials before automatically granting credit to participants. University participant pools provide a unique opportunity for crowdsourcing on a minimal budget because participants are compensated with course credit rather than money.

More recently, crowdsourcing has been available via commercial platforms, such as the Amazon Mechanical Turk, or MTurk, platform^2^. MTurk is a popular venue for crowdsourcing due to the large number of available workers and the relative ease with which tasks can be uploaded and payments disbursed. Methods for crowdsourcing and estimates of data quality have been available for years, and several recommendations have emerged from past work. For example, collecting multiple responses per image can account for natural variation and the relative skill of the untrained workers (23). Furthermore, a majority vote of MTurk workers can label images with similar accuracy to that of experts (19). Although those studies were limited to labeling categorical features of stock images, other studies have shown success with more complex stimuli. For example, MTurk workers were able to diagnose disease and identify the clinically relevant areas in images of human retinas with accuracy approaching that of medical experts (17). Amazon’s MTurk is a particularly valuable tool for researchers because it provides incentives for high quality work. The offering party has the ability to restrict their task to only workers with a particular work history, or a more general criterion known as ‘Master Turk’ status. The Master title is a status given to workers by Amazon based on a set of criteria that Amazon believes to represent the overall quality of the worker; note that Amazon does not disclose those criteria.

The time and cost savings of using crowdsourcing to label data are obvious, but crowdsourcing is only a viable solution if the output is sufficiently accurate. The goal of the current project was to test whether crowdsourcing image labels (also called tags) could yield a sufficient positive-data training set for ML from image-based phenotypes in as little as a single day. We focus on corn tassels for this effort but we anticipate our findings to extend to other similar tasks in plant phenotyping.

In this project, we recruited three groups of people for our crowdsourcing tassel identification task, from the two online platforms Sona and MTurk. The first group consisted of students recruited for course credit, or the Course Credit group. The second group consisted of paid Master-status Mechanical Turk workers, (the Master MTurkers group), and the third group consisted of paid non-master Mechanical Turk workers (the non-Master MTurkers group). The accuracy of the different groups’ tassel identification was evaluated against an expert-generated gold standard. These crowdsourced labeled images were then used as training data for a “bag-of-features” machine learning algorithm.

We found that performance of Master and non-Master MTurkers was not significantly different; however both groups performed better than the Course Credit group. At the same time, using the labeling data from either course credit, MTurk or Master MTurk did not make any significant difference in the performance of the machine learning algorithm when trained on sets generated by any of these groups. We conclude that crowdsourcing via MTurk can be useful for establishing ground truth sets for complex image analysis tasks in a short amount of time, and that MTurkers’ and expert MTurkers’ performance exceed that of students working for course credit. At the same time, perhaps surprisingly, we also show that the differences in labeling quality do not significantly affect the performance of a machine learning algorithm trained by any of the three groups.

## Methods

### Ethics Statement

Research involving human participants was approved by the Institutional Review Board at Iowa State University under protocol 15-653.

### General outline

The overall scheme of the work is depicted in Figure 1. Course Credit, Master MTurkers, MTurkers, and an expert, all labeled corn tassels in a set of 80 images. First, the labeling performance was assessed against the gold standard. Then, each set of labeled images was also used to train a bag-of-features machine learning method. The trained methods were each tested against a separate expert-labeled training set, to assess how differently the ML method performed with different training sets.

**Figure 1:**
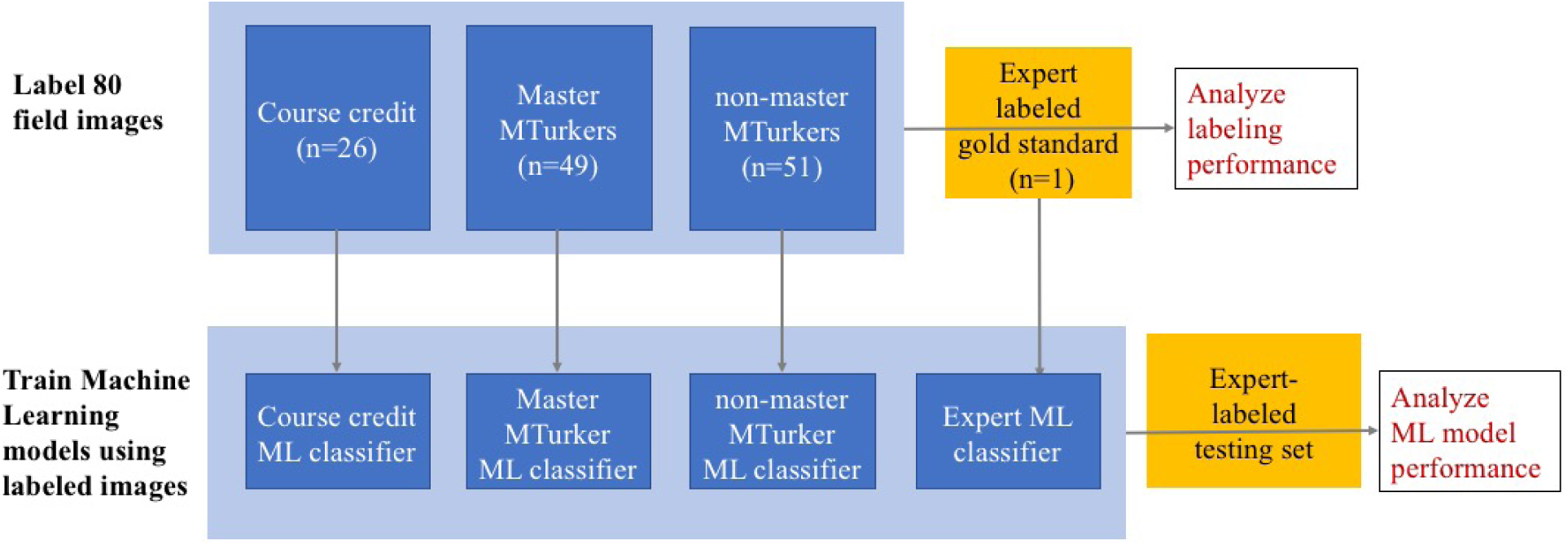
Overall schema of datasets (boxes) and processes (arrows) that led to the analyses (red). **Top row:** The Expert Labeled dataset was used a gold standard to analyze how well the different experimental groups (blue boxes) performed. **Bottom row:** the labeling from each experimental group was used to train an ML classifier. Each ML classifier was then tested against an expert-labeled test set.

### Recruiting Participants

The Course Credit group included 30 participants, which were recruited using the subject pool software Sona from the undergraduate psychology participant pool at Iowa State University. Recruited students were compensated with course credits. The master MTurkers included 65 master-qualified workers recruited through MTurk. The exact qualifications for master status are not published by Amazon, but are known to include work experience and employer ratings of completed work. Master MTurkers were paid $8.00 to complete the task and the total cost was $572.00. Finally, the non-master MTurkers pool included 66 workers with no qualification restriction, recruited through the Amazon Mechanical Turk website. Due to the nature of Amazon’s MTurk system, it is not possible to recruit only participants who are *not* master qualified. However, the purpose of including the non-master MTurkers was to evaluate workers recruited without the additional fee imposed by Amazon for recruitment of Masters MTurkers. Non-master MTurkers were also paid $8.00 to complete the task and the total cost was $568.00. Note that the costs include Amazon’s fees. Of the 30 students recruited, 26 completed all 80 images. Of the 65 Master MTurkers recruited, 49 completed all images. Of the 66 non-master MTurkers recruited, 51 completed all images. Data collected from participants who did not complete the survey were not included in subsequent analyses.

### Pilot Study

A brief cropping task was initially administered to Sona and master MTurkers groups as a pilot study to test the viability of this project and task instructions. Each participant was presented with a participant-specific set of 40 images randomly chosen from 393 total images. The accuracy of participant labels helped designate Easy and Hard status for each image. Forty images were classified as “easy to crop”, and 40 as “hard to crop”, based on accuracy results of the pilot study. An expert who made gold standard boxes made adjustments to the Easy/Hard classifications based on personal experience. These 80 images were selected for the main study. As opposed to the pilot study, participants in the main study each received the same set of 80 images, with image order randomized separately for each participant. The results of the pilot study indicated that at least 40 images could be processed without evidence of fatigue so the number of images included in the main experiment was increased to 80. The pilot study also indicated, via user feedback, that a compensation rate of $8.00 for the set of 80 images was acceptable to the MTurk participants. To expedite the pilot study, we did not include regular MTurkers. Our rationale was that feasibility for a larger study could be assessed by including master MTurkers and Sona only.

### Gold Standard

We define a *gold standard box* for a given tassel as the box with the smallest area among all bounding boxes that contain the entire tassel, a minimum bounding box. Gold-standard boxes were generated by the expert, a trained and experienced researcher. The expert cropped all 80 images then computationally minimized the boxes to be minimum bounding. These images were used to evaluate the labeling performance of crowdsourced workers, and should not be confused with the ‘ground truth’ which were used to refer the labeled boxes used in training the ML model.

### General Procedure

We selected the images randomly from a large image pool obtained as part of an ongoing maize phenomics project. The field images focused on a single row of corn captured by cameras set up as part of the field phenotyping of the maize Nested Association Mapping (28), using 456 cameras simultaneously, each camera imaging a set of 6 plants. Each camera took an image every 10 minutes during a two week growing period in August 2015 (15). Some image features varied, for example, due to weather conditions and visibility of corn stalks, but the tassels were always clearly visible. Images were presented on a Qualtrics webpage^3^ and Javascript was used to provide tassel annotation functionality. After providing Informed Consent, participants viewed a single page with instructions detailing how to identify corn tassels and how to create a minimum bounding box around each tassel. Participants were first shown an example image with the tassels correctly bounded with boxes (Figure 2). Below the example, participants read instructions on how to create, modify, and delete bounding boxes using the mouse. These instructions explained that an ideal bounding box should contain the entire tassel with as little additional image detail as possible. Additional instructions indicated that overlapping boxes and boxes containing other objects would sometimes be necessary and were acceptable as long as each box accurately encompassed the target tassel. Participants were also instructed to only consider tassels in the foreground, ignoring tassels that appear in the background. After reading instructions, participants clicked to progress to the actual data collection. No further feedback or training were provided. The exact instructions are provided in the Supplementary Materials.

**Figure 2:**
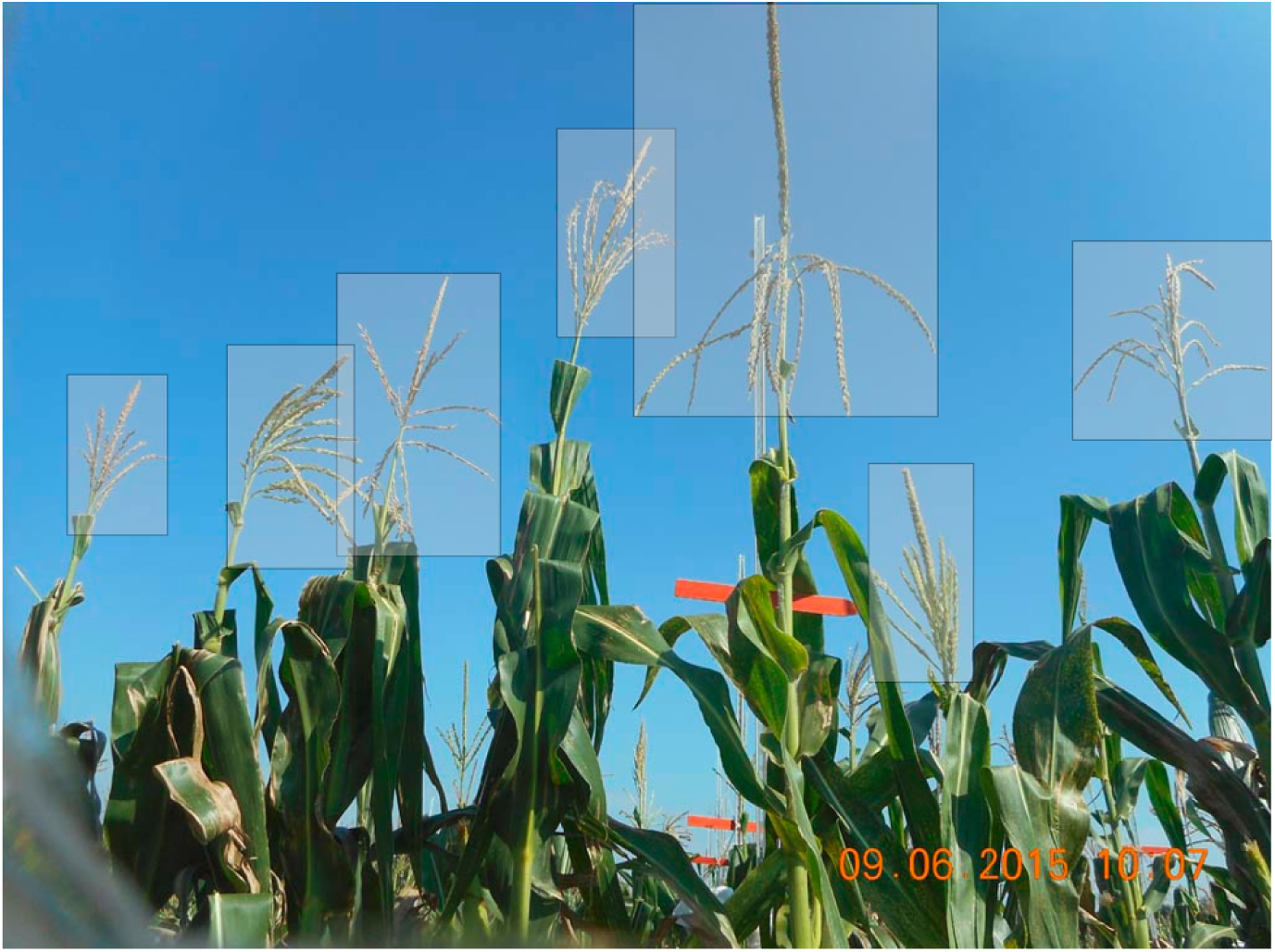
Example image used during training to demonstrate correct placement of bounding boxes around tassels

For each image, participants created a unique bounding box for each tassel by clicking and dragging the cursor. Participants could subsequently adjust the vertical or horizontal size of any drawn box by clicking on and dragging a box corner, and could adjust the position of any drawn box by clicking and dragging in the box body. Participants were required to place at least one box on each image before moving on to the next image. No upper limit was placed on the number of boxes. Returning to previous images was not allowed. The time required to complete each image was recorded in addition to the locations and dimensions of user-drawn boxes.

### Defining Precision and Recall

Consider any given participant-drawn box and gold standard box as in the right panel of Figure 3. Let *PB* be the area of the participant box, let *GB* be the area of the gold standard box, and let *IB* be the area of the intersection between the participant box and the gold standard box. Precision (*Pr*) is defined as *IB/PB*, and recall (*Rc*) is defined as *IB/GB*. Both *Pr* and *Rc* range from a minimum value of 0 (when the participant box and gold standard box fail to overlap) to a maximum value of 1 (full overlap of boxes). As an overall measure of performance for a participant box as an approximation to a gold standard box, we use *F*1, the harmonic mean of precision and recall:

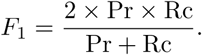

**Figure 3:**
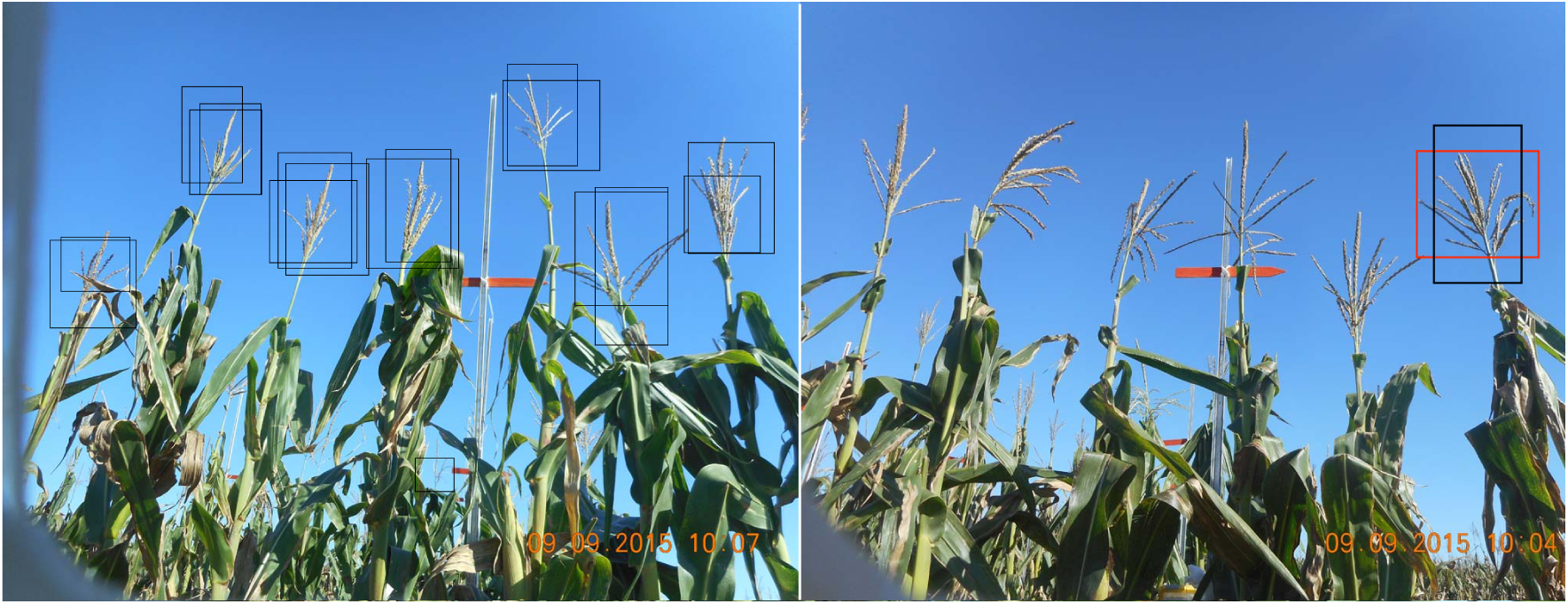
Left: Sample participant-drawn boxes. Right: The Red box is the gold standard box and black is a participant-drawn box

Each participant-drawn box was matched to the gold standard box that maximized *F*1 across all gold standard boxes within the image containing the participant box. If more than one participant box was matched to the same gold standard box, the participant box with the highest *F*1 value was assigned the *Pr, Rc*, and *F*1 values for that match, and the other participant boxes matching that same gold standard box were assigned *Pr, Rc*, and *F*1 values of zero. In the usual case of a one-to-one matching between participant boxes and gold standard boxes, each participant box was assigned the *Pr, Rc*, and *F*1 values associated with its matched gold standard box.

To summarize the performance of a participant on a particular image, *F*1 values across participant-drawn boxes were averaged to obtain a measure referred to as *F*_*mean*_. This provides a dataset with one performance measurement for each combination of participant and image that we use for subsequent statistical analysis.

## Results

### Data Distribution

As described above, precision and recall were calculated for each participant-drawn box. Density of precision recall pairs by group based on 61,888 participant-drawn boxes are shown in the heatmap visualization of Figures 4a,4b and 4c.

**Figure 4:**
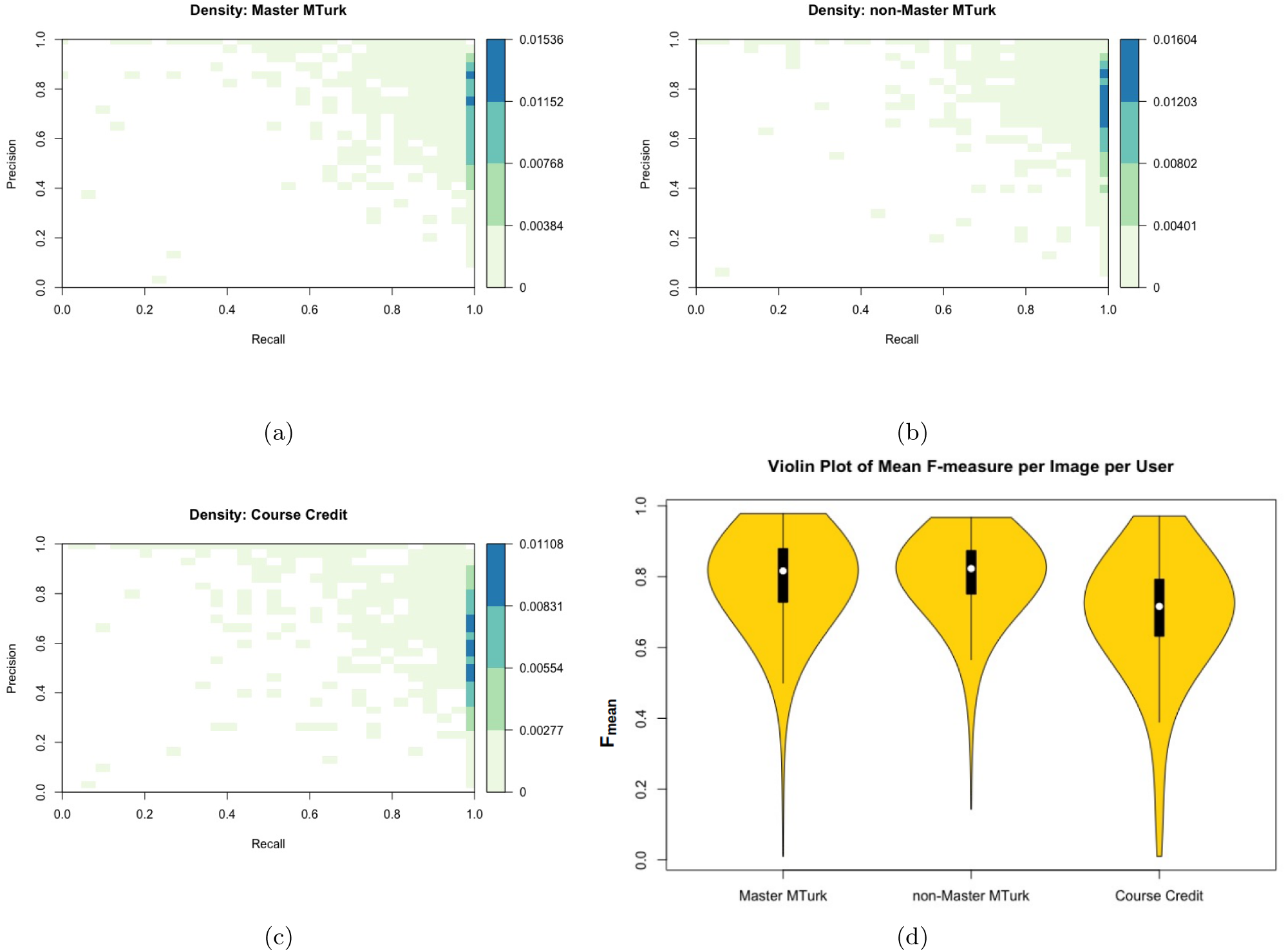
Density of precision recall pairs by group based on 61,888 participant-drawn boxes for **a**: Master MTurkers, **b**: MTurkers, and **c**: Course Credit participants. **d**: violin plots showing the distribution of F-measure per image per user, where white circles: distribution median; black bars: second and third quartiles; black lines 95% confidence intervals.

High value precision-recall pairs are more common than low value precision-recall pairs in all three groups. Perfect recall values were especially common because participants tended to draw boxes that encompassed the minimum bounding box, presumably to ensure that the entire tassel was covered.

### Testing for Performance Differences among Groups

Figure 4d shows the distribution of *F*_*mean*_ for the three groups. We used a linear mixed-effects model analysis to test for performance differences among groups with the *F*_*mean*_ value computed for each combination of image and user as the response variable. The model included fixed effects for groups (Master MTurker, non-Master MTurker, course credit), random effects for participants nested within groups, and random effects for images. The *mixed* procedure available in SAS software was used to perform this analysis with the Kenward-Roger method (11) for computing standard errors and denominator degrees of freedom. The analysis shows significant evidence for differences among groups (*p*-value < 0.0001). Furthermore, pairwise comparisons between groups (Table 1) show that both Master and non-Master MTurkers performed significantly better than undergraduate students performing the task for course credit. There was no significant performance difference between Master and non-Master MTurkers.

**Table 1:** Parameter estimates from the ANOVA with master MTurk group as baseline.

|  | Estimate | Standard Error | p-value |
| --- | --- | --- | --- |
| non-master MTurk vs. Master MTurker | 0.01125 | 0.02078 | 0.5893 |
| Course Credit vs. Master MTurker | -0.1005 | 0.02521 | 0.0001 |
| Course Credit vs. non-master MTurk | -0.1117 | 0.01517 | $<0.0001$ |

### Time Usage and Fatigue

Next we wanted to understand whether there is any change of time taken to annotate over the task given, whether there is a significant difference between the groups, and specifically if any change indicated fatigue. Participants took a median time of 26.43 seconds to complete an image, with the median time for the Master MTurker group at 30.02 seconds, non-Master MTurkers at 29.40 seconds, and the course credit student group at 16.86 seconds. The course credit group generally spent less time than either MTurker group. It is worth noting that there is a large variance in time spent on each image, with the longest time for a single image at 15,484.63 seconds, and the shortest being 0.88 seconds. The very long image annotation time was probably due to the participant taking a break after cropping part of the image and then coming back later to finish that image.

There is a general downward trend in the time spent on each image over time. The trend is shown in Figure 5a, via linear regression on log time with fixed effects for group, question ordinal index and group × question ordinal index, and random effects for user and image. The trend is statistically significant in all three groups, with similar effect sizes. As participants complete questions, the average time spent per question is reduced by about 1%, as shown by Table 2. By looking at the interaction term between participant group and question index, we were able to conclude that the reduced time effect is not significantly different between the Master MTurker and non-Master MTurker group (p=0.6003), but is different between the course credit group and Master MTurker group (p=0.0431). This difference is weakened in terms of course credit versus non-Master MTurker, with a p-value of 0.1086.

**Table 2:** Parameter estimates in linear mixed effects regression of time spent each image

| | Estimate ($\hat{\beta}$) | Exponential of Estimate ($\exp(\hat{\beta})$) | p-value |
| --- | --- | --- | --- |
| Master MTurk | -0.01043 | 0.9896 | $< 0.0001$ |
| non-Master MTurk | -0.01073 | 0.9893 | $< 0.0001$ |
| Course Credit | -0.01181 | 0.9883 | $< 0.0001$ |

**Figure 5:**
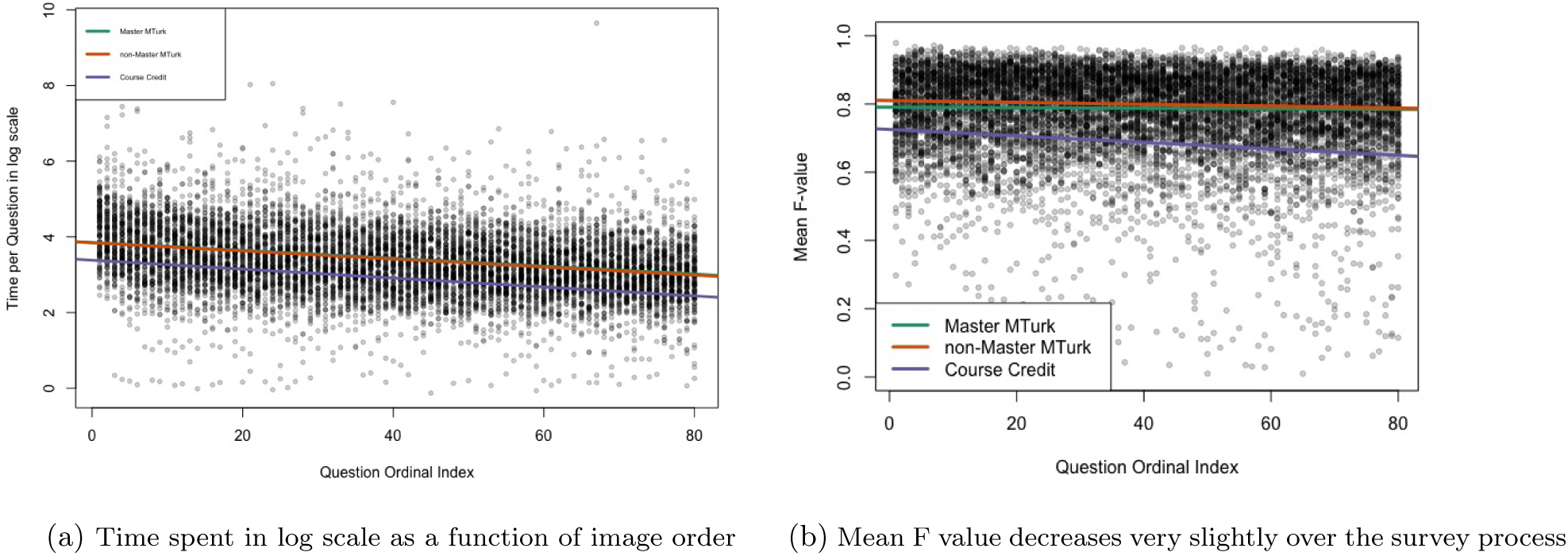
Accuracy metrics as a function of progression through the task.

We also analyzed the change in accuracy, as measured by *F*_*mean*_ as the test progresses. Figure 5b shows that *F*_*mean*_ decreased slightly as the task progressed. The decreases are statistically significant (p<0.05) for all three groups. However, the effect sizes (average decrease in *F*_*mean*_ per round of image) for both MTurker groups are almost negligible, with Master MTurk group showing a 0.00008 decrease per image and Non-master group showing a 0.00027 decrease. Decrease in *F*_*mean*_ for the course credit group is only slightly more noticeable, at 0.00095, and on a scale of 0-1 is unlikely to affect training. To summarize the effect of image order, there was a subtle decline in *F*_*mean*_ and a larger decrease in image completion time as the survey progressed.

Another question of interest was whether image accuracy correlates with image completion time. Indeed, there tended to be a slight increase in accuracy as time spent on an image increased. Although the correlation is statistically significant in all three groups, the effect sizes are too small to conclude that spending more time on a single image has an important positive effect on accuracy for that image.

In conclusion, all three groups of participants spent less time on each image as the survey progressed, possibly due to increasing familiarity in the task. Although their performance in the task also decreases slightly over time, the effects were almost negligible. This fatigue effect, while significant, is minor.

### Image Difficulty

Did the annotators spend more time on more difficult images? To answer this question, we obtained the Best Linear Unbiased Predictor (BLUP) (8) of each image in the above analyses to assess whether each image contributes to increased or decreased accuracy and time. BLUPs can be viewed as predictions of random effects, in our case, one prediction of the eighty images. Figure 6 is a scatter plot, with each point representing an image. The horizontal axis shows the BLUPs with regard to logtime. The higher the BLUP, the more this particular image contributes to increased time spent on each question. Similarly, the vertical axis shows the BLUPs with regard to *F*_*mean*_. Images with higher BLUPs tended to be processed more accurately. We also obtained a difficult / easy classification of all eighty images from our expert who manually curated the gold standard boxes, as they are shown by the two different colors on the plot.

**Figure 6:**
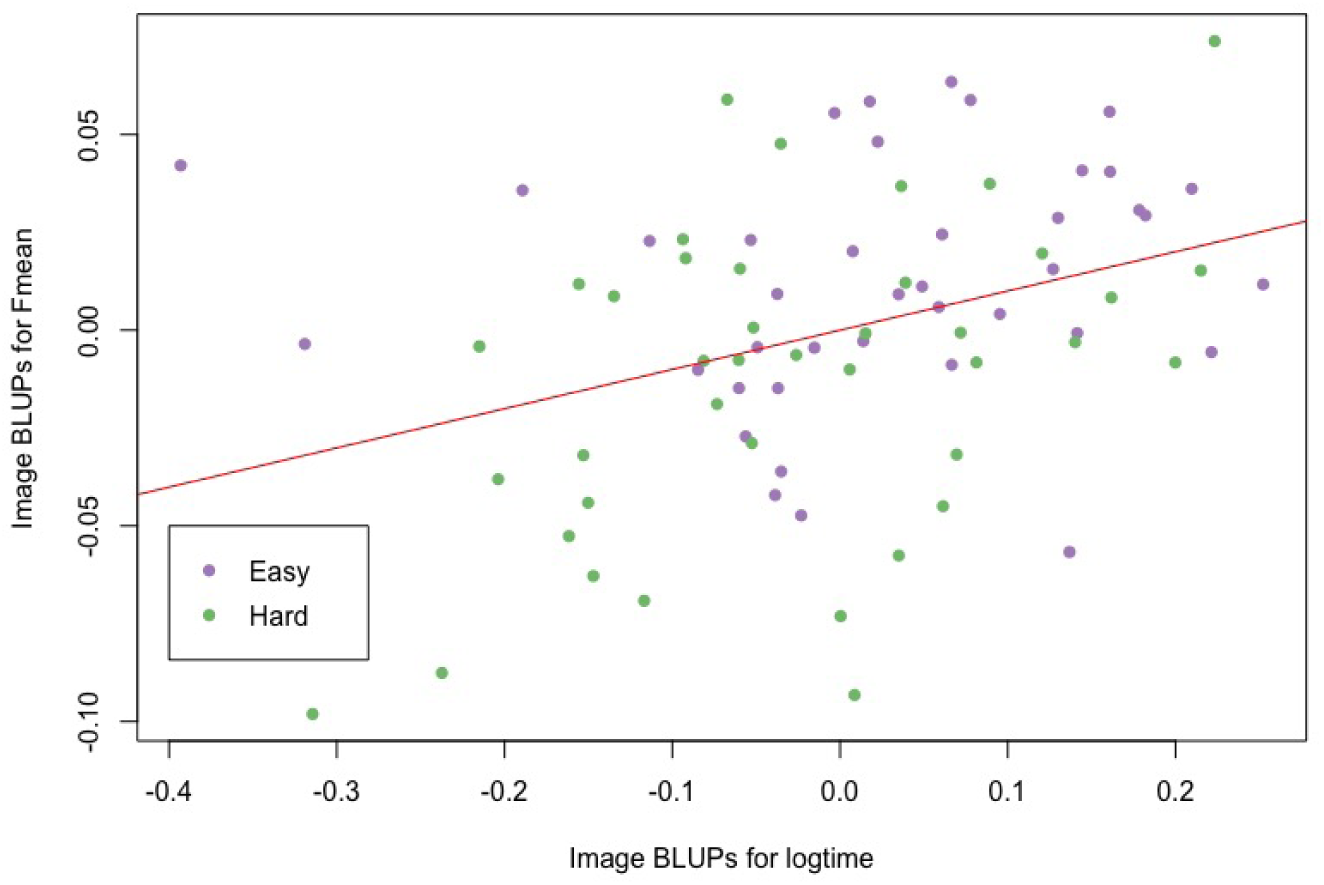
Best Linear Unbiased Predictors for image in analyses for *F*_*mean*_ and time in log scale. Color represents image difficulty determined by expert.

It is interesting to observe that longer time spent annotating an image correlates positively with accuracy. Indeed, the linear regression fit shown as the red line on the plot has a slope estimate of 0.1003 (*p*=0.00136), and an adjusted *R*_2_ of 0.1127, suggesting weak correlation. Furthermore, the images that our expert con-sidered difficult did not take participants longer to complete, nor did they yield significantly lower accuracy. The images were shown to participants in a random order, eliminating the possibility that fatigue contributes to the longer time it takes to complete easy images. Since previous analysis showed that participants tend to spend less time on images shown to them later (Figure 5a), this may suggest ordering the images so that more difficult images are shown to the participants first. In that way, a surveyor may take advantage of the fact that participants tend to spend more time on each image in the beginning, to obtain more accurate results.

### Evaluating the Performance of Participant-Trained Classifiers

Automatic tassel detection is an important prerequisite for fast computation of quantitative traits. We can automatically detect tassels in images using a classifier trained with data derived from crowdsourcing. Although our results above show that paid Master MTurkers and non-Master MTurkers tend to provide higher quality tassel bounding boxes than students working for course credit, the differences in quality we have detected may not necessarily translate into better training of a classification algorithm. We therefore examined how the performance of a classification algorithm varies as the data used to train the classifier varies across participants.

The algorithm consists of two stages. The first stage involves extracting features from a set of training images using a bag-of-features method (18). Each training image corresponds to a single box within one of our original images. The training images (i.e., boxes) are selected so that each contains either a tassel or no tassel. Each training image is then represented by a vector of frequencies, with one frequency for each feature. The second stage of the algorithm involves training a support vector machines (SVM) classifier using the frequency vectors associated with the training images, as well as the status of each training image: whether it contains a tassel or not. For each of the 126 participants in our study, we constructed a set of training images using the participant-drawn boxes as the positive set (i.e., the training images containing a tassel), together with a constant negative set of 600 images (corresponding to boxes that contain no tassel). The number of training images in the positive set varied across participant, with a median of 457. The classification algorithm was then separately trained using each of the 126 training datasets.

We applied the 126 participant-trained classifiers to a test set of 600 tassel images and 600 non-tassel images. Performance was calculated as the mean of the true positive rate and the true negative rate of the classification. Overall, the algorithm achieved a classification performance of 0.8811, averaged over all participants. For the master and non-master MTurker groups, the average performances were 0.8851 and 0.8781, respectively. For the course credit group, it was 0.8795. We performed a linear model analysis of the 126 performance measures to test for group differences. The F test yielded a p-value of 0.7325, indicating no detectable differences among the average performances of the three groups.

We also trained a classifier based on the ground truth data that the our expert curated. This classifier achieved an accuracy of 0.91, slightly higher than the performance of the classifiers trained from the crowdsourced labels.

## Discussion

Machine learning methods have revolutionized processing and extracting information from images, and are being used in fields as diverse as public safely, biomedicine, weather, military, entertainment, and, in our case, agriculture. However, these algorithms still require an initial training set created by expert individuals before structures can be automatically extracted from the image and labeled. This project has identified crowdsourcing as a viable method for creating initial training sets without the time consuming and costly work of an expert. Our results show that straightforward tasks, such tassel cropping, do not benefit from the extra fee assessed to hire master over non-master MTurkers. Performance between the two groups was not significantly different, and non-master MTurkers can safely be hired without compromising data quality.

The MTurk platform allows for fast collection of data within a day instead of one to two weeks. While MTurk may be one of the most popular crowdsourcing platforms, many universities possess a research participant pool that compensates students with class credit instead of cash for their work. However, in our study the undergraduate student participant pool did not perform as well as either of the MTurker groups. While it is possible that MTurk workers are simply more conscientious than college students, it is also possible that monetary compensation is a better motivator than course credit. In addition to the direct monetary reward, both groups of MTurkers were also motivated by either working towards or maintaining the “master” status. Such implicit motivational mechanisms might be useful in setting up a long-term crowdsourcing platform. The distinction in labeling performance between MTurkers and students does not matter when considering the actual outcome of interest: how well the machine learning algorithm identifies corn tassels when supplied with each of the three training sets. The accuracy of the ML algorithm used here was not affected by the quality of the training set provided, which were manually-labeled through crowdsourcing. Therefore, a student participant pool with a non-monetary rewards system provides the opportunity for an alternate model by lowering overall image tagging cost. This would allow additional features to be tagged or a larger number of responses to be sourced with existing funding levels and further database expansion.

Indeed, there are many crowdsourcing projects that do not offer monetary reward. For example, the Backyard Worlds: Planet 9 project hosted by NASA for search of planets and star systems in space (12), the *Phylo* ^4^ game for multiple sequence alignment (10) and *fold.it* ^5^ (4) for protein folding. These projects attract participants by offering the chance to contribute to real scientific research. This concept has been categorized as citizen science, where nonprofessional scientists participate in crowdsourced research efforts. In addition to the attraction of the subject matter, these projects often have interactive and entertaining interfaces to quickly engage the participants’ interests and attention. Some of them were even designed as games, and competition mechanisms such as rankings provide extra motivation. Another important purpose of such citizen science projects is to educate the public about the subject matter. Given the current climate regarding Genetically Modified Organisms (GMOs), crowdsourcing efforts of crop phenomic and phenotypic research could potentially be a gateway to a better understanding of plant research in the general public. A recent effort has shown that non-experts can be used for accurate image-based plant phenomics annotation tasks (6). However, the current data points to the challenge of non-monetary reward in sustaining a large-scale annotation effort.

Phenomics is concerned with the quantitative and qualitative study of phenomes, where all possible traits of a given organism vary in response to genetic mutations and environmental influences (9). An important field of research in phenomics is the development of high-throughput technology analogous to high-throughput sequencing in genetics and genomic studies, to enable the collection of large-scale data with minimal efforts. Many phenotypic traits could be recorded with images, and databases such as BioDIG (20) make the connection of such image data with genomic information, providing genetics researchers with tools to examine the relationship between the two types of data directly. Hence, the computation and manipulation of such phenomic image data becomes essential. In plant biology, maize is central for both basic biological research as well as crop production (reviewed in (14)). As such, phenotypic information derived from ear (female flowers) and tassel (male flowers) are key to both the study of genetics and crop productivity: flowers are where meiosis and fertilization occur as well as the source of grain. To add a new features such as tassel emergence, size, branch number, branch angle and anthesis to the systems such as BioDIG, the specific tassel location and structure should be located, and our solution to this task is to use crowdsourcing combined with machine learning to reduce cost and time of such a pipeline, while expanding its utility. Our findings, and the suggested crowdsourcing methods can be generally applied to other phenomic analysis tasks. It is worthy to note that differences in quality of training sets may not translate into significant differences in classification, as was in our study. However, this may vary between different classification algorithms, and different training sets. We hope our study will help establish some best practices for researchers in setting up such a crowdsourcing study. Given the ease and relatively low cost of obtaining data through Amazon’s Mechanical Turk, we recommend it over the undergraduate research pool. That being said, student research pools would be a suitable method for obtaining proof of concept or pilot data to support a grant proposal.

## Acknowledgements

These will be added after manuscript is accepted.

## Data and Software

The software for this project is available from: https://github.com/ashleyzhou972/Crowdsource-Corn-Tassels

The data for this project are available from: https://doi.org/10.6084/m9.figshare.6360236.v2

## Footnotes

1 www.sona-systems.com

2 https://www.mturk.com/

3 www.qualtrics.com

